# Menstrual cycle selectively elevates anticipatory effort cost without altering reward sensitivity in human motivation

**DOI:** 10.64898/2026.06.05.730335

**Authors:** Nitsan Schwarz, Daniel Harlev, Noham Wolpe

## Abstract

Motivation fluctuates across the menstrual cycle, yet the computational mechanisms underlying these changes remain unclear. We tested whether hormonally defined cycle phases selectively alter distinct components of effort-based motivation in naturally cycling women (n=51), who completed an effort-reward decision-making task and an effort psychophysics task in both the late-follicular and mid-luteal phases, alongside electrocardiography and ecological momentary assessment. Hierarchical Bayesian modelling revealed that the anticipated cost of physical effort (effort sensitivity) was selectively elevated in the mid-luteal phase, with no corresponding change in reward sensitivity. The luteal increase in effort sensitivity was attenuated in women who entered that phase after days of higher affective valence and arousal, indicating that positive affective state buffers cyclical motivational vulnerability. Complementing these findings, phase-related individual differences in effort the mapping between objective and perceived effort (effort differentiation) were not different by phase, but were moderated by heart rate variability and momentary affective states, revealing stable person-level variation in how the cycle shapes effortful experience. Together, these results identify effort sensitivity as a specific computational mechanism of cyclical motivational change, with implications for understanding the elevated burden of cycle-related psychiatric conditions across the female reproductive lifespan.

## INTRODUCTION

Psychological fluctuations in mood, irritability and motivation are among the most common and disruptive experiences reported by women across the reproductive lifespan^1^. Yet despite their prevalence and link to vulnerability to mental health conditions^2^, the mechanisms through which the menstrual cycle shapes mental health remains poorly understood. So far, women’s reproductive physiology remains a notable blind spot in cognitive and computational neuroscience.

A central challenge in understanding cycle-linked motivational changes is that motivation itself is not a unitary construct. Recent advances in computational neuroscience have shown that motivated behaviour can be decomposed into dissociable components, each of which can be altered independently^3^. When choosing whether to act, individuals weigh both the subjective value of anticipated rewards (measured by reward sensitivity), and the subjective cost of the effort required to obtain them (measured by effort sensitivity)^3,4^. These two parameters can be estimated separately from effort-reward decision-making tasks and have been shown to dissociate in clinical populations. Motivational impairment in depression, schizophrenia, and apathy syndromes is driven predominantly by elevated effort sensitivity, with some evidence for blunted reward sensitivity^3,5^. Asking how the menstrual cycle affects motivation therefore requires asking which of these components is altered: whether cyclical motivation reflects changes in what women value, in what they are willing to do to obtain it, or in both.

Motivation also unfolds across two temporal stages that are themselves distinguishable^6^. While decision-making captures the anticipatory weighting of effort and reward before an action is taken, a separate experiential component can be captured during the action itself^6^. Effort perception is typically measured using self-report methods which are susceptible to biases, e.g., in self-report anchoring^7^. Effort differentiation, however, quantifies the mapping between subjective rating and objective effort, which overcomes many of these reporting biases. Reduced effort differentiation is associated with depressive symptoms in the typical adult population^8^. As anticipatory and experiential effort processing are dissociable^9^ and rely on partially distinct neural substrates^10^, either could plausibly be affected by cyclical hormonal state. A full characterisation of how the menstrual cycle shapes motivation and effort processing, therefore, requires measuring both.

How might hormonal and physiological changes across the menstrual cycle affect anticipatory and experiential effort? Oestradiol and progesterone modulate similar dopaminergic and GABAergic circuits that underlie effort-based decision-making^11,12^. Oestradiol broadly facilitates dopaminergic neurotransmission^13^ and progesterone acts through its metabolite allopregnanolone on GABA-A receptors^14^. In terms of effort perception, changes in autonomic regulation, indexed by changes in heart rate variability (HRV) across the menstrual cycle^15^, can influence interoceptive and emotional processing linked to effort perception^16^.

Here, we asked how the menstrual cycle shapes motivation and effort processing across both its anticipatory and experiential components: Naturally cycling women completed an effort-reward decision-making task and an effort psychophysics task in both the mid-luteal and late-follicular phases of the menstrual cycle, with phases verified through both retrospective and physiological measurements. In-session echocardiographic assessment of HRV and twice-daily ecological momentary assessments of affective state across a full cycle were acquired. We preregistered that reward sensitivity, effort sensitivity, and effort differentiation would be higher in the late-follicular than mid-luteal phase. We further preregistered moderation analyses testing whether phase-related effects would vary as a function of HRV, particularly increased HRV in the late-follicular phase. Finally, we preregistered analyses testing whether ecological momentary assessments of emotional dimensions of valence and arousal in the three days preceding each session would predict individual differences in effort and reward processing across phases.

## METHODS

All primary hypotheses, including power analyses and analysis plan for the two behavioural tasks were preregistered at: https://aspredicted.org/32gz-pw5p.pdf and https://aspredicted.org/33m9rz.pdf. Additional analyses are labelled as exploratory.

### Participants

Participants were healthy, naturally cycling women aged 18-35 years. Inclusion criteria was based on gold-standard menstrual cycle research^17^. We recruited female participants who reported a regular menstrual cycle between length of 22-35 days, no use of hormonal contraception or hormone-altering medication, no pregnancy or breastfeeding, and no psychiatric medication use. Participants were recruited via university posters, social media advertisements, and word-of-mouth. All participants provided written informed consent prior to participation, and the study was approved by the Tel Aviv University Research Ethics Committee (ref: 0008220-2). Each laboratory session took approximately 60 minutes, and participants were compensated for their time.

### Study overview

A within-subject design was used in which each participant completed two laboratory sessions scheduled in distinct menstrual cycle phases, alongside ecological momentary assessment (EMA) collected continuously across one menstrual cycle (Fig. 1). Each laboratory session followed the same structure. Participants first underwent maximal voluntary contraction (MVC) calibration using a hand-held isometric dynamometer. The calibration defined a session-specific scaling of force, such that all subsequent effort demands were expressed relative to the participant’s current force capacity. Participants then completed an effort psychophysics (EP) task followed by an effort-reward decision-making (DM) task probing choice under varying reward and force (effort) requirements. Electrocardiography (ECG) was recorded prior to task participation during rest and during MVC calibration.

**Figure 1.**
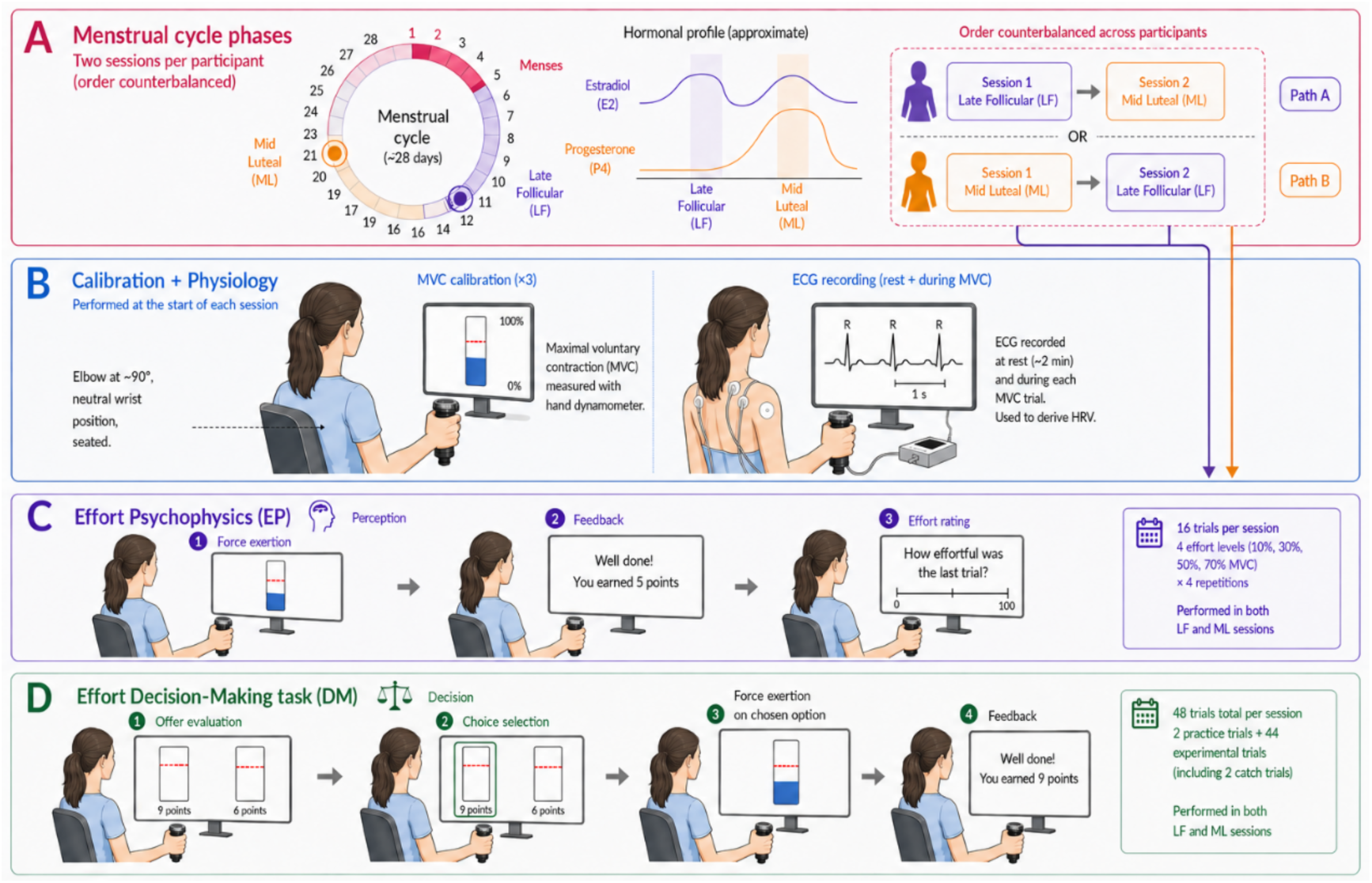
Overall study design. (A) Menstrual-cycle sampling scheme. Each participant completed two laboratory sessions in a counterbalanced order: one session in the late-follicular (LF) phase and one session in the mid-luteal (ML) phase. Shaded regions indicate the approximate hormonal profiles of oestradiol and progesterone across the cycle; arrows indicate the counterbalanced session order across participants. (B) Calibration and physiology procedures performed at the beginning of each session. Maximal voluntary contraction (MVC) was measured with a hand dynamometer to calibrate task effort levels. Electrocardiogram (ECG) activity was recorded continuously throughout the session, including during a resting baseline period and during MVC trials, to derive heart-rate variability (HRV) measures. (C) Effort psychophysics (EP) task. In each trial, participants exerted force at a target level and then rated the subjective effort of the just-completed trial on a 0-100 scale. Each session included 16 trials comprising four effort levels (10%, 30%, 50%, and 70% MVC), with four repetitions per level. (D) Effort decision-making (DM) task. In each trial, participants evaluated two effort-reward options, selected one option, exerted the required force on the chosen option, and then received performance feedback. Each session included 44 experimental trials (2 catch trials), preceded by 2 practice trials.

All analyses were performed in R version 4.32^18^ and MATLAB version R2023b. All force-based tasks were performed using a hand-held dynamometer (BIOPAC, USA; TSD121C grip force transducer). Force signals were sampled at 250 Hz. Visual feedback, task presentation and behavioural recording were implemented using a custom experimental interface programmed in Python with PsychoPy library ^19^. ECG was recorded continuously using a standard three-electrode configuration connected to the BIOPAC ECG100C amplifier module. Signals were sampled at 1000 Hz and were recorded using BIOPAC Student Lab software.

At the beginning of each laboratory session, participants completed an MVC calibration procedure consisting of three maximal squeezes of the dynamometer. The first trial was performed without visual feedback. The second and third trials were performed with real-time visual feedback displayed as a vertical force bar on the screen. The display included current force output, the maximum achieved force, and a reference marker corresponding to 110% of the previous maximum in order to encourage maximal exertion. We have tested the reliability of this procedure in an independent sample of participants on repeated measurements, which showed excellent reliability (Supplementary Results, Fig. S1). The highest force recorded across the three trials was defined as the MVC for that session. All subsequent force requirements in both EP and DM tasks were expressed as percentages of this session-specific MVC.

### Determining menstrual cycle phase and time of in-lab testing

Cycle phase was determined using a combination of retrospective tracking and physiological verification, as defined in standard menstrual cycle studies^17^. Participants tracked their menstrual cycles for a minimum of two months prior to study initiation to ensure cycle regularity. To index ovulation, participants were instructed to perform daily self-administered luteinising hormone testing (Life, Israel) beginning two days prior to the expected ovulation and for a maximum of five days. When a positive luteinising hormone signal was obtained, ovulation was defined relative to that event. When no positive test was recorded, ovulation timing was estimated retrospectively using backward counting from the onset of the subsequent menstrual period, in line with standard procedures in menstrual cycle studies^17^.

To capture the late-follicular and mid-luteal phases, laboratory sessions were scheduled for −4 to 0 days relative to ovulation (late-follicular), −9 to −5 days relative to next menses (mid-luteal)^17^. After data collection, phase assignments were validated using all available cycle information. Analyses requiring a direct comparison between menstrual phases were restricted to participants who contributed one physiologically and retrospectively verified late-follicular session and one retrospectively verified mid-luteal session. In total, 7 participants were excluded due to phase misalignment, leaving a total of 44 participants for the phase comparisons.

### Effort-reward decision-making task

An effort-reward decision-making task assessed anticipatory effort valuation, adapted from previous work^20^. Each session included a practice block followed by 44 experimental trials and 2 catch trials embedded within the trial set. Catch trials were defined as trials in which one option strictly dominated the other by offering both higher reward and lower effort. Catch-trial performance was used to confirm task comprehension; no participant met the preregistered exclusion criterion of failing both catch trials within a session.

On each trial, participants were presented with two options differing in reward magnitude and required effort. Reward ranged from 1 to 10 points, and effort ranged from 10% to 90% of MVC. Effort/reward option combinations were chosen so as to maximise parameter recovery (see Supplementary Results). Option positions (left/right) were randomised across trials. Participants selected an option using the computer mouse. Following selection, they exerted the required force using the dynamometer. Force execution followed the same success criteria used in the EP task. Feedback indicated success or failure and the reward obtained.

Decision-making behaviour was modelled using a hierarchical Bayesian framework, as done before ^20^. For each option, subjective value was defined as:

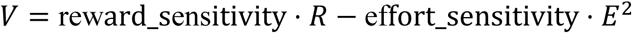

where 𝑅 is reward magnitude and 𝐸is effort level as proportion MVC. Effort is squared as done in previous research^20,21^. Choice probabilities were modelled as:

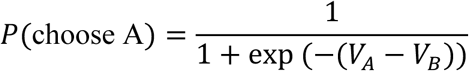

The model was fitted using Markov chain Monte Carlo sampling. Convergence was assessed using standard diagnostics, including visual inspection of trace plots and R-hat values. Model fit was further evaluated using posterior predictive checks. The primary decision-making outcomes were effort sensitivity and reward sensitivity. Phase effects on these parameters were tested in the phase-verified sample by comparing late-follicular and mid-luteal sessions.

### Effort psychophysics task

The EP task was designed to quantify the mapping between subjective effort experience and objective force required^8^. On each trial, participants viewed a vertical thermometer display with a fixed target line and were instructed to squeeze the dynamometer to raise a bar to reach the target. Critically, the visual target remained constant across trials, whereas the mapping between applied force and bar movement was manipulated by varying the gain between force input and visual output. This required participants to exert different force levels to achieve the same visual target.

Four force levels were used, corresponding to 10%, 30%, 50%, and 70% of MVC. Each level was repeated four times, yielding 16 experimental trials following 2 practice trials. Trials were presented in pseudo-random order. A trial was classified as successful if the participant reached and maintained the required force level for a cumulative duration of 3 seconds within a 5-second response window. Following each trial, participants received feedback indicating success or failure and then rated perceived effort on a visual analogue scale ranging from 0 to 100, with anchors stating ‘no effort at all’ for 0 and ‘maximum effort’ for 100.

The preregistered primary EP outcome was effort differentiation, operationalized as the within-session linear slope relating subjective effort rating to objective force level. A steeper slope indicates greater perceptual differentiation between low and high force demands, whereas a flatter slope indicates reduced differentiation^8^. The preregistered phase comparison tested whether effort differentiation differed between late-follicular and mid-luteal sessions. In accordance with the preregistration, participants were excluded from the primary phase-based EP analysis if they did not contribute a retrospectively verified late-follicular and mid-luteal session pair or if their force-rating Spearman correlation deviated by more than 2.5 standard deviations from the sample mean. In total, 10 participants were excluded as a result of these criteria, leaving a sample of 41.

### Moderators of task-phase relationship

In addition to the main effect of menstrual phase on the primary measures of decision-making (effort and reward sensitivity) and EP (effort differentiation) tasks, we examined additional moderators of the task-phase relationship. Specifically, we examined whether trait mental health questionnaires, physiological (HRV) and emotional state (EMA) measures moderated the task-phase relationship.

For assessing mental health traits, participants completed self-report questionnaires assessing depressive symptoms, anxiety symptoms, apathy, and menstrual-cycle-related symptom burden. These included the Patient Health Questionnaire-9 (PHQ-9^22^), the Generalized Anxiety Disorder-7 (GAD-7^23^), the Apathy Motivation Index (AMI^24^), and the Daily Record of Severity of Problems (DRSP^25^). Trait-like questionnaire measures were summarised descriptively across the sample, whereas DRSP was additionally examined in the preregistered phase-sensitive analyses.

To assess physiology and the autonomic nervous system, ECG was acquired at rest and during an active task. ECG was first measured continuously for five minutes while participants were sat at rest. Active ECG was then recorded for the duration of the MVC task (∼3 minutes). ECG signals were processed offline in R, following standard recommendations for short-term HRV analysis^26^. Signals were band-pass filtered (0.5-40 Hz), R-peaks were detected using a prominence-based algorithm, and RR intervals were extracted and cleaned prior to HRV estimation. HRV was quantified as the root mean square of successive differences (RMSSD), a standard time-domain index of short-term vagally mediated heart rate variability ^27^:

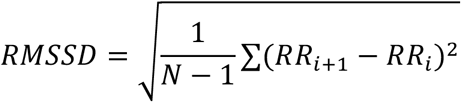

RMSSD was derived separately for rest (baseline) and MVC (active) conditions. Because the MVC segment was substantially shorter than the baseline recording, separate quality-control thresholds were applied for the two conditions. Sessions were retained for baseline HRV analyses only if at least 30 cleaned RR intervals were available, and for active MVC HRV analyses only if at least 10 cleaned RR intervals were available. This approach was chosen because RMSSD is relatively robust in ultra-short recordings ^28^. Sessions were excluded if ECG acquisition failed, if too few R-peaks were detected, or if the cleaned RR count fell below these predefined thresholds. HRV predictors were standardized prior to entry into the statistical models.

To assess participants’ emotional state outside the in-lab sessions, participants completed twice-daily EMA assessments for a total of one continuous menstrual cycle (cycle length mean=28.83 days, SD=2.81 days). Each assessment included visual-analogue scale ratings of mood, anxiety, energy, and fatigue, followed by a working memory task and a perceived effort rating measurement, as done in our previous research^29^. Arousal was defined as energy plus the inverse of fatigue, and valence was defined as mood (sad-happy) plus anxiety (anxious-calm)^29^. For predicting in-lab session-level variables from EMAs, we examined mean and variability of arousal and valence 3 days preceding each laboratory session, as preregistered. For linking EMA to EP, mean arousal, arousal variability, mean valence, and valence variability were entered as session-level moderators of the trial-level force-rating relationship in the mixed-effects models described above. For the effort-based decision-making task, the same pre-session EMA variables were tested as predictors of model-derived effort sensitivity and reward sensitivity.

## RESULTS

### Participant characteristics and data inclusion

Participant characteristics and raw descriptive measures by menstrual-cycle phase are presented in Table 1. Of the 51 naturally cycling participants with full dataset recruited, seven participants were excluded as their phase could not be retrospectively or physiologically validated as preregistered: four participants contributed two luteal-type sessions and therefore could not contribute a follicular-versus-luteal comparison, whereas three additional participants contributed sessions retrospectively classified as ovulatory rather than late-follicular. Separate analyses on an independent sample confirmed high test-retest reliability of MVC procedure across repeated laboratory sessions (Supplementary Results and Fig. S1).

**Table 1.**
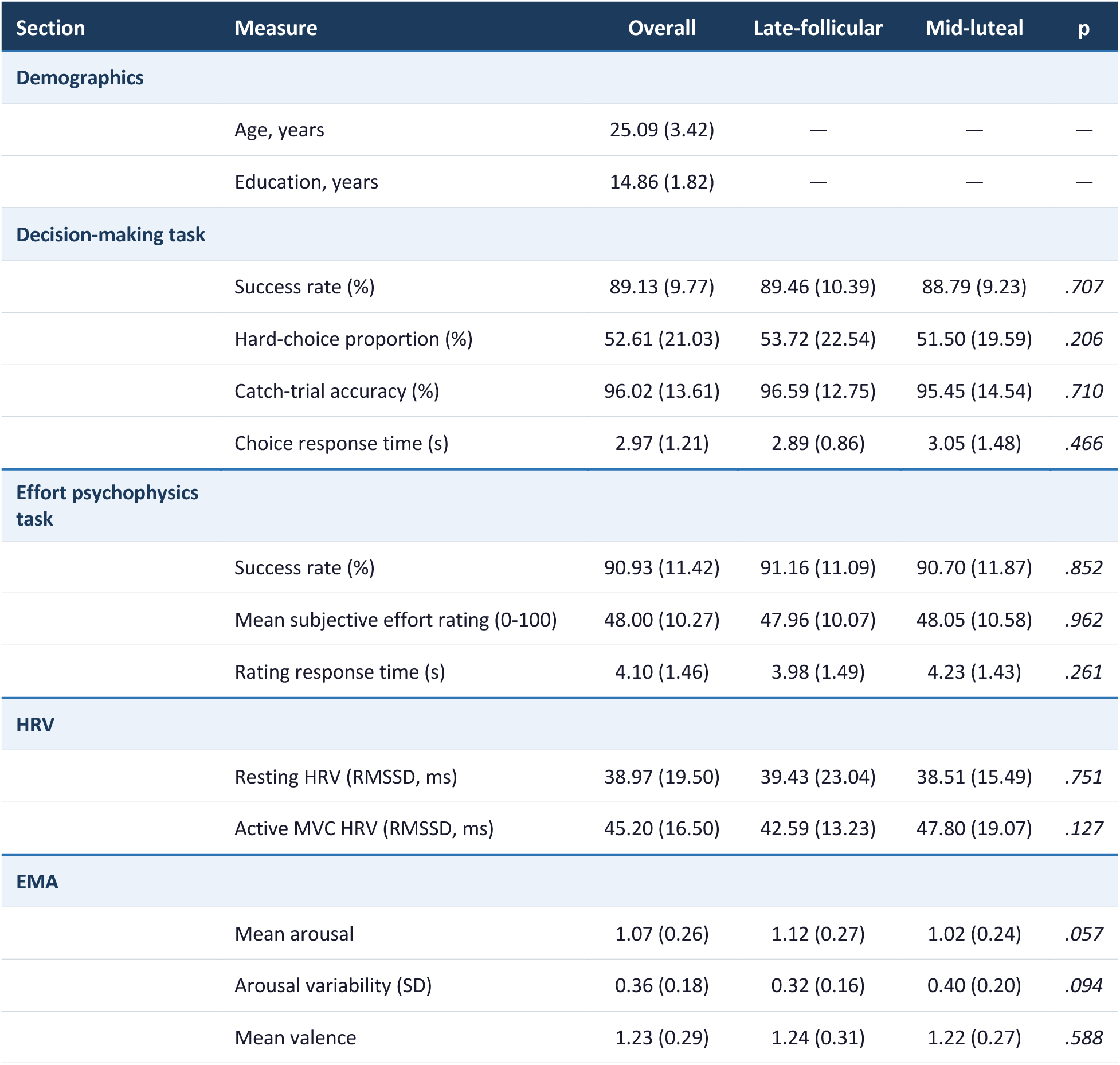
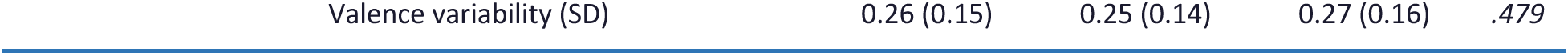
Sample characteristics and raw descriptive measures by menstrual-cycle phase. Values are mean (SD). Phase-specific columns show descriptives for the late-follicular and mid-luteal sessions. *P*-values reflect paired-samples comparisons between phases. HRV = heart rate variability; RMSSD = root mean square of successive differences; MVC = maximal voluntary contraction; EMA = ecological momentary assessment. Dashes (—) indicate measures not collected by phase.

| Section | Measure | Overall | Late-follicular | Mid-luteal | p |
| --- | --- | --- | --- | --- | --- |
| Demographics |  |  |  |  |  |
|  | Age, years | 25.09 (3.42) | — | — | — |
|  | Education, years | 14.86 (1.82) | — | — | — |
| Decision-making task |  |  |  |  |  |
|  | Success rate (%) | 89.13 (9.77) | 89.46 (10.39) | 88.79 (9.23) | .707 |
|  | Hard-choice proportion (%) | 52.61 (21.03) | 53.72 (22.54) | 51.50 (19.59) | .206 |
|  | Catch-trial accuracy (%) | 96.02 (13.61) | 96.59 (12.75) | 95.45 (14.54) | .710 |
|  | Choice response time (s) | 2.97 (1.21) | 2.89 (0.86) | 3.05 (1.48) | .466 |
| Effort psychophysics task |  |  |  |  |  |
|  | Success rate (%) | 90.93 (11.42) | 91.16 (11.09) | 90.70 (11.87) | .852 |
|  | Mean subjective effort rating (0-100) | 48.00 (10.27) | 47.96 (10.07) | 48.05 (10.58) | .962 |
|  | Rating response time (s) | 4.10 (1.46) | 3.98 (1.49) | 4.23 (1.43) | .261 |
| HRV |  |  |  |  |  |
|  | Resting HRV (RMSSD, ms) | 38.97 (19.50) | 39.43 (23.04) | 38.51 (15.49) | .751 |
|  | Active MVC HRV (RMSSD, ms) | 45.20 (16.50) | 42.59 (13.23) | 47.80 (19.07) | .127 |
| EMA |  |  |  |  |  |
|  | Mean arousal | 1.07 (0.26) | 1.12 (0.27) | 1.02 (0.24) | .057 |
|  | Arousal variability (SD) | 0.36 (0.18) | 0.32 (0.16) | 0.40 (0.20) | .094 |
|  | Mean valence | 1.23 (0.29) | 1.24 (0.31) | 1.22 (0.27) | .588 |
|  | Valence variability (SD) | 0.26 (0.15) | 0.25 (0.14) | 0.27 (0.16) | .479 |

### Effort and reward sensitivity in decision-making

The decision-making analyses were conducted on the full phase-verified sample of 44 participants. Catch-trial performance indicated good task comprehension overall: across the 88 subject-by-phase sessions, 81 sessions contained no catch-trial errors and seven sessions contained one error, and no participant met the preregistered exclusion criterion of failing both catch trials within a session.

Across the reward-effort space, phase differences in hard-choice behaviour were not uniformly distributed, but instead clustered primarily at higher effort levels, with comparatively smaller variation along the reward dimension (Fig. 2A). This pattern suggests that the phase effect in the task was driven more strongly by effort-cost weighting than by reward valuation. At the aggregate behavioural level, however, basic task performance was similar across phases (full descriptive statistics are reported in the Supplementary Results and Fig. S2).

**Figure 2.**
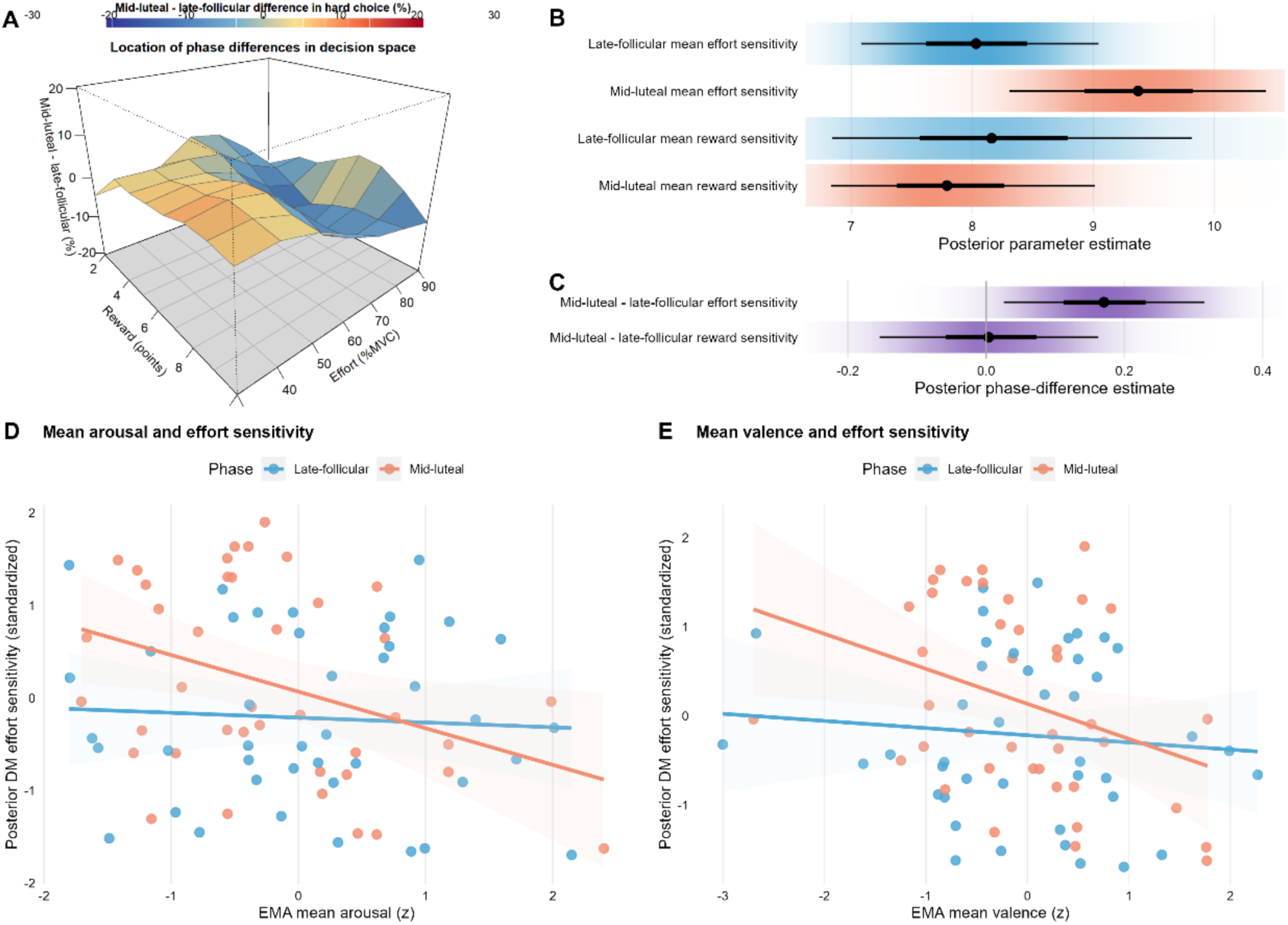
Decision-making task phase effects, decision space differences, and EMA associations. (A) Phase differences in decision space. The surface shows the mid-luteal minus late-follicular difference in hard-choice proportion across the offered reward-effort space. Positive values (warmer colours) indicate a higher proportion of hard choices in the mid-luteal phase; negative values (cooler colours) indicate a higher proportion in the late-follicular phase. For visualisation purposes, the displayed surface was smoothed by interpolation across the grid. (B) Posterior mean parameter estimates by phase. Blue and orange rows show the late-follicular and mid-luteal phase-specific mean estimates for effort sensitivity and reward sensitivity, respectively. Points indicate posterior means and horizontal bars indicate 95% credible intervals. (C) Posterior phase-difference estimates. Purple rows show the posterior mid-luteal minus late-follicular contrasts for effort sensitivity and reward sensitivity. Points indicate posterior means and horizontal bars indicate 95% credible intervals. (D, E) Associations between EMA mean arousal (D) and mean valence (E) and standardised effort sensitivity, shown separately for the late-follicular and mid-luteal phases. Points represent participant-phase observations, lines show phase-specific linear fits, and shaded bands indicate 95% confidence intervals.

The primary hierarchical Bayesian modelling analyses focused on the model-derived parameters of effort sensitivity and reward sensitivity. Effort sensitivity showed a clear phase effect. Mean effort sensitivity was 8.03 (SD = 3.96) in the late-follicular phase and 9.39 (SD = 4.40) in the mid-luteal phase (Fig. 2B), and the estimated population-level mid-luteal minus late-follicular difference was 0.171, 95% CrI [0.006, 0.350], with a posterior probability of 0.978 (Fig. 2C). By contrast, reward sensitivity did not vary reliably by phase. Mean reward sensitivity was 8.20 (SD = 6.06) in the late-follicular phase and 7.84 (SD = 4.55) in the mid-luteal phase (Fig. 2B), and the estimated mid-luteal minus late-follicular difference was 0.004, 95% CrI [-0.185, 0.196], with a posterior probability of .513 (Fig. 2C). Thus, menstrual phase selectively modulated the weighting of effort costs during decision-making, increasing effort sensitivity by 19% at the group level in the mid-luteal relative to late-follicular phase, whereas reward sensitivity remained stable.

We next tested whether questionnaire (DRSP), HRV, and EMA variables moderated these decision-making parameters. DRSP and HRV did not show reliable phase-dependent associations with either effort sensitivity or reward sensitivity (full results reported in the Supplementary Results). By contrast, emotional state prior to the in-lab session, namely arousal and valence averaged three days prior to the session both showed phase-dependent associations with effort sensitivity. Mean arousal interacted significantly with phase (beta = 0.044, SE = 0.017, *t* = 2.63, *df* = 38.30, *p* = .012, 95% CI [0.010, 0.077]), and mean valence showed a similar interaction (beta = 0.031, SE = 0.015, *t* = 2.04, *df* = 38.15, *p* = .048, 95% CI [0.0003, 0.063]). Follow-up simple-slope analyses indicated that in the mid-luteal phase, higher mean arousal and higher mean valence were more strongly associated with lower effort sensitivity, whereas the corresponding late-follicular relationships were flatter (Fig. 2D-E). No comparable EMA effects were observed for reward sensitivity. Exploratory AMI analyses pointed in the same motivational direction, such that higher AMI was associated with greater effort sensitivity, but not reward sensitivity. (non-phase-based results are reported in full in the Supplementary Results and Fig. S3).

### Perceptual effort differentiation

The effort psychophysics analyses were conducted on 41 participants, as three participants were excluded because their rating-force non-parametric correlation deviated by more than 2.5 standard deviations from the sample mean, as preregistered.

Subjective effort ratings increased reliably as required force increased, confirming that participants differentiated between the four force levels during the task (beta = 0.867, SE = 0.019, *t* = 45.11, *df* = 1245.34, *p* < .001; Fig. 3A). Success rates were high and did not differ between phases, with means of 91.16% (SD = 11.09%) in the late-follicular phase and 90.70% (SD = 11.87%) in the mid-luteal phase (*t*(40) = 0.19, *p* = .852, 95% CI [-4.46, 5.38]). Rating response times were likewise comparable across phases, with means of 3.98 s (SD = 1.49) and 4.23 s (SD = 1.43), respectively (*t*(40) = -1.14, *p* = .261, 95% CI [-0.69, 0.19]).

**Figure 3.**
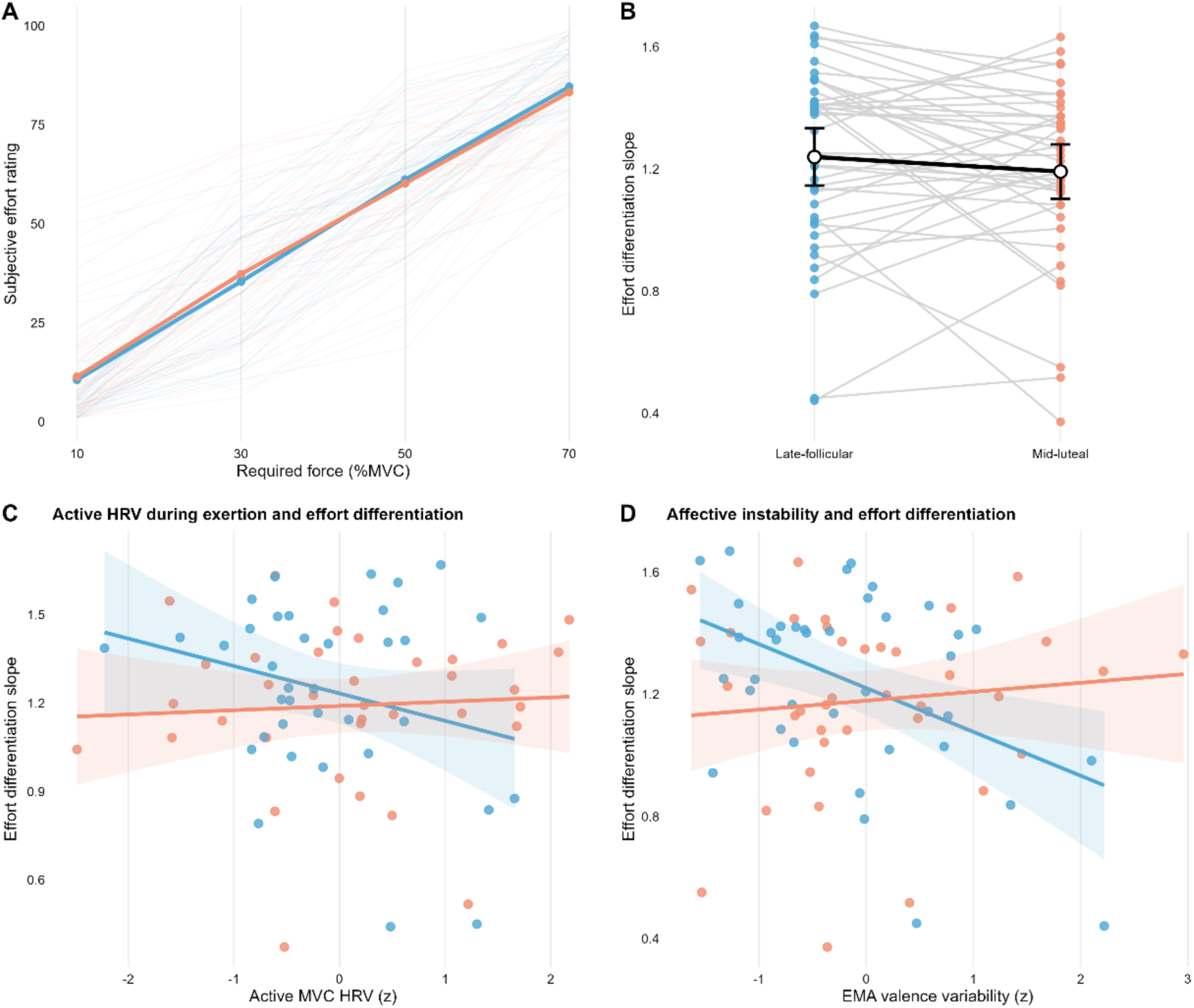
Effort psychophysics task performance, phase comparison, and moderators of effort differentiation. (A) Mean subjective effort rating as a function of required force level (%MVC) in the late-follicular and mid-luteal phases. Thin lines show participant-level mean rating functions within each phase, whereas thick lines and points show phase-level means at each force level. (B) Effort differentiation slope by phase. Effort differentiation was operationalised as each participant’s within-session linear slope relating subjective effort ratings to objective force level. Points show individual participants, connecting lines indicate within-participant pairing across phases, and black summary markers indicate phase means and 95% confidence intervals. (C) Association between heart rate variability recorded during MVC trials (active HRV) and effort differentiation slope, shown separately for the late-follicular and mid-luteal phases. (D) Association between EMA valence variability and effort differentiation slope, shown separately for the late-follicular and mid-luteal phases. In panels C and D, points represent participant-phase observations, lines show phase-specific linear fits, and shaded bands indicate 95% confidence intervals.

The primary analysis tested whether effort differentiation differed between the late-follicular and mid-luteal phases. Contrary to the preregistered hypothesis, effort differentiation did not differ significantly between phases, with a mean follicular-minus-luteal difference of 0.048 (*t*(40) = 1.12, *p* = .269, 95% CI [-0.038, 0.133]) (Fig. 3B). Thus, menstrual phase did not produce a reliable shift in perceptual differentiation of increasing physical effort at the group level.

We next examined possible moderations of questionnaire, physiological (HRV), and EMA on rating-force relationship. The DRSP did not significantly moderate effort differentiation, as the force x DRSP interaction was not significant (beta = -0.019, SE = 0.019, *t* = -1.01, *df* = 1245.34, *p* = .312, 95% CI [-0.057, 0.018]), nor was the force x phase x DRSP interaction significant (beta = 0.023, SE = 0.027, *t* = 0.86, *df* = 1245.34, *p* = .392, 95% CI [-0.029, 0.075]).

By contrast, HRV showed a more selective pattern. Resting HRV showed a trend-level association with flatter force-rating slopes (beta = -0.035, SE = 0.018, *t* = - 1.95, *df* = 1119.91, *p* = .051, 95% CI [-0.070, 0.000]), but this association did not differ by phase (beta = 0.001, SE = 0.028, *t* = 0.04, *df* = 1119.91, *p* = .972, 95% CI [-0.054, 0.056]). However, higher active HRV was associated with flatter rating-force slopes (beta = -0.064, SE = 0.023, *t* = -2.77, *df* = 1121.67, *p* = .006, 95% CI [-0.110, - 0.019]), and this association differed by phase (beta = 0.084, SE = 0.029, *t* = 2.91, *df* = 1121.67, *p* = .004, 95% CI [0.027, 0.141]). These results indicate that higher active HRV was associated with flatter effort differentiation in the late-follicular phase, whereas this negative association was attenuated in the mid-luteal phase (Fig. 3C).

EMA measures derived from the 3 days preceding each laboratory session also showed a selective pattern. Mean arousal showed a trend-level positive association with effort differentiation (beta = 0.036, SE = 0.018, *t* = 1.93, *df* = 1166.86, *p* = .054, 95% CI [-0.001, 0.072]), but did not interact with phase (beta = 0.029, SE = 0.029, *t* = 0.99, *df* = 1166.86, *p* = .323, 95% CI [-0.028, 0.086]). Greater variability in valence, however, was associated with flatter force-rating slopes (beta = -0.098, SE = 0.021, *t* = -4.65, *df* = 1150.94, *p* < .001, 95% CI [-0.139, -0.057]), and this association differed significantly by phase (beta = 0.119, SE = 0.029, *t* = 4.16, *df* = 1150.94, *p* < .001, 95% CI [0.063, 0.176]). This pattern indicates that the link between greater valence instability and poorer effort differentiation was strongest in the late-follicular phase and attenuated in the mid-luteal phase (Fig. 3D). The remaining non-significant effort differentiation moderator results are reported in the Supplementary Results.

Exploratory AMI analyses pointed in the same direction as the moderator results: higher AMI was associated with reduced effort differentiation, but these did not depend on phase (full results in the Supplementary Results and Fig. S3).

## DISCUSSION

Across the reproductive lifespan, women commonly report that energy, drive, and motivation fluctuate with the menstrual cycle^30^. While motivation has been discussed in context of hormonal fluctuations^30^, what has been missing up until now is a specification of *which* element of motivation such symptoms reflect. Here we show that the cycle acts selectively: in the mid-luteal phase, women weighted the anticipated cost of effort approximately 19% more heavily, without affecting how they value reward nor how they perceive effort. Cyclical motivational symptoms are therefore better understood not as a loss of interest in rewards, nor as a global sense that everything feels harder; but as a specific, anticipatory increase in the cost of action.

This selective effect of cycle phase on anticipatory effort cost appears to be at odds with the body of literature focussing on reward processing across the menstrual cycle. Neuroimaging and behavioural work have linked the high oestradiol follicular phase to stronger reward-related responses and reward sensitivity, with attenuation in the luteal phase^31,32^. In the present study, however, cycle phase was associated with altered effort sensitivity rather than reward sensitivity. This is notable because elevated effort costs are increasingly recognised as a transdiagnostic feature of motivational impairment^4,33^. These findings therefore suggest that the luteal phase may transiently increase the subjective cost assigned to effort, even in healthy naturally cycling women. More speculatively, such a mechanism could help explain why the luteal phase is a period of heightened psychiatric vulnerability and symptom exacerbation in some individuals^1^: rather than generating motivational symptoms de novo, cyclical hormonal changes may periodically increase effort costs in those already close to a clinical threshold.

The direction of this effect was opposite to our preregistered prediction, which expected effort sensitivity to be higher in the high-oestradiol late-follicular phase. The observed luteal increase is therefore less consistent with the oestradiol-dopamine account that motivated our hypothesis, and may instead implicate mechanisms associated with the progesterone-dominant luteal phase. Oestradiol has been linked to facilitation of mesolimbic dopaminergic transmission^13,34^, a system implicated in lowering the subjective cost of effort^35^. By contrast, the luteal phase is characterised by increases in progesterone and its metabolite allopregnanolone, which may dampen phasic mesolimbic dopamine signalling^36,37^. A luteal increase in effort sensitivity could therefore reflect reduced dopaminergic support for effortful action during this phase. However, this interpretation remains speculative, as most direct evidence coming from animal models is mixed, showing no effect of the intact cycle on effort-based choice and hormone effects that were dose-dependent and sometimes opposing^11^. No human study, to our knowledge, has directly linked luteal neurosteroids to effort-based choice, which remains a question for future research.

The same effort-cost computation was sensitive to motivational vulnerability across multiple timescales. Trait apathy, measured using the AMI^24^, independently predicted higher effort sensitivity, consistent with work in non-clinical samples^38^. Recent affective context also mattered: participants who entered the luteal phase after days of higher mean valence and arousal showed a smaller increase in effort sensitivity. This finding suggests that affective state before the luteal window may shape vulnerability within it, consistent with computational accounts in which affect modulates effort-based choice^1,39^. More broadly, the results position effort-based decision-making as a marker not only of stable motivational traits, but also of dynamic motivational state^39^. The convergence of trait apathy, recent affect, and hormonal phase on the same effort-cost parameter raises the possibility that premenstrual motivational vulnerability is modifiable, with wellbeing in the preceding days buffering subsequent effort costs. By contrast, affective variability, rather than mean affect, predicted experiential effort differentiation, suggesting a potentially distinct pathway linking affective instability to the subjective experience of effort.

While effort sensitivity rose in the mid-luteal phase, perceived effort differentiation did not shift at the group level, reinforcing the view that the cycle alters the anticipatory valuation of effort rather than its experienced intensity during action. Where perception did vary, it tracked autonomic engagement rather than phase: in the follicular phase, women with *higher* ‘active’, task-engaged HRV showed flatter force-rating slopes, discriminating less sharply between effort levels. Because effortful action typically elicits vagal withdrawal as resources are mobilised and reduced HRV^40^, the higher HRV *during* exertion plausibly indexes weaker engagement with the task in these individuals. It may thus be this disengaged state, not poorer regulatory capacity, that blunts effort discrimination. This suggestion is consistent with neurovisceral integration accounts in which dynamic vagal control, rather than resting tone, couples interoceptive and central processes^41^. That the effect was specific to the follicular phase suggests cycle physiology gates how strongly autonomic engagement shapes effort perception; a pattern which resting HRV would miss. Taken together, our findings situate effort processing as a candidate mechanism for cyclical mental-health vulnerability. Premenstrual worsening has been reported across several psychiatric conditions, including depression, anxiety disorders, and bipolar disorder^1,42^, all of which commonly involve motivational difficulty^6^. Our results implicate anticipatory effort sensitivity, rather than the experiential perception of effort, as a potential computational pathway through which cyclical physiology shapes motivational state. As a task-based and state-sensitive measure^39^, effort sensitivity may provide information that is not captured by retrospective symptom ratings, which remain vulnerable to recall bias, stigma, and limited introspective access to the processes governing choice^43^. This framework generates a clear clinical prediction: PMDD should be associated with exaggerated luteal increases in effort sensitivity relative to typical cyclers, a possibility that remains to be tested in clinical populations.

This framework has implications for intervention. Cognitive behavioural therapy reduces premenstrual symptom burden^44,45^, but the mechanisms through which behavioural treatments improve cyclical symptoms remain poorly specified. Our finding that pre-luteal affective state buffered the luteal increase in effort sensitivity suggests one candidate pathway: interventions that improve affective context before the luteal window may attenuate subsequent effort-cost amplification. This possibility is supported by evidence that cognitive-behavioural components can shift effort-based decision-making parameters in healthy participants^20^, indicating that effort-cost computations are modifiable. Thus, alongside previously established evidence for altered reward-circuit responses in PMDD, including reduced ventral-striatal responses to positive stimuli^46^, our results point to anticipatory effort valuation as a mechanistically specific and potentially modifiable component of cyclical motivational symptoms.

A key strength of our study is that it separated components of motivation that are often conflated. By combining an effort-reward decision-making task with an effort psychophysics task, we could distinguish anticipatory effort valuation from reward valuation and from the subjective perception of effort during action^17^. Concurrent HRV and EMA measures further allowed us to situate these computational parameters within autonomic and affective contexts, linking laboratory-derived measures of motivation to dynamic physiological and experiential states outside the lab. Moreover, the within-participant design and rigorous phase assignment procedure further strengthen inference relative to studies relying on between-participant comparisons or retrospective cycle counting alone. Several limitations nevertheless constrain interpretation. Most importantly, we did not include hormonal assays, meaning that phase effects cannot be directly attributed to measured concentrations of oestradiol, progesterone, or their metabolites. The task also focused on physical effort; whether similar phase-related changes occur for cognitive effort, which may be especially relevant to everyday motivational symptoms^47^, remains unknown. Finally, the observed direction of the effort-sensitivity effect, with higher effort sensitivity in the luteal than follicular phase, was opposite to our preregistered prediction. This mismatch should be considered in light of the limited evidence base available for generating directional hypotheses in women’s health studies, particularly the scarcity of computational psychiatry studies in women’s reproductive health.

In conclusion, women’s reproductive physiology remains a major blind spot in computational and clinical neuroscience, and cyclical motivational symptoms have lacked a mechanistic account proportionate to their prevalence and burden. Our findings begin to provide such an account: across the menstrual cycle, motivational change reflected a selective, state-dependent elevation in the anticipated cost of effort during the luteal phase. By separating effort cost from reward value and perceived exertion within the same participants, we show that the menstrual cycle can act on one component of motivation while leaving others comparatively unchanged. Moving towards mechanism-informed assessment is therefore a necessary step for characterising women’s cyclical mental health, and may ultimately improve detection and treatment of vulnerability to mental health conditions across the reproductive lifespan.

## ACKNOWLEDGEMENTS

We thank the participants for their engagement and participation in this study.

## SUPPLEMENTARY MATERIAL

### Supplementary Results

We tested the reliability of our maximal voluntary contract (MVC) assessment in an independent sample. The independent sample comprised 56 participants (female: *n* = 40), with a mean age of 27.38 years (SD = 5.83; range: 19-59 years). The majority of participants were right-handed (*n* = 53; left-handed: *n* = 3). Test-retest reliability of MVC measurements was assessed using a two-way mixed-effects model for absolute agreement, treating participants as a random factor and sessions as a fixed factor. Reliability was excellent, ICC(2,1) = 0.948, based on 227 observations across participants (Fig. S1).

**Figure S1.**
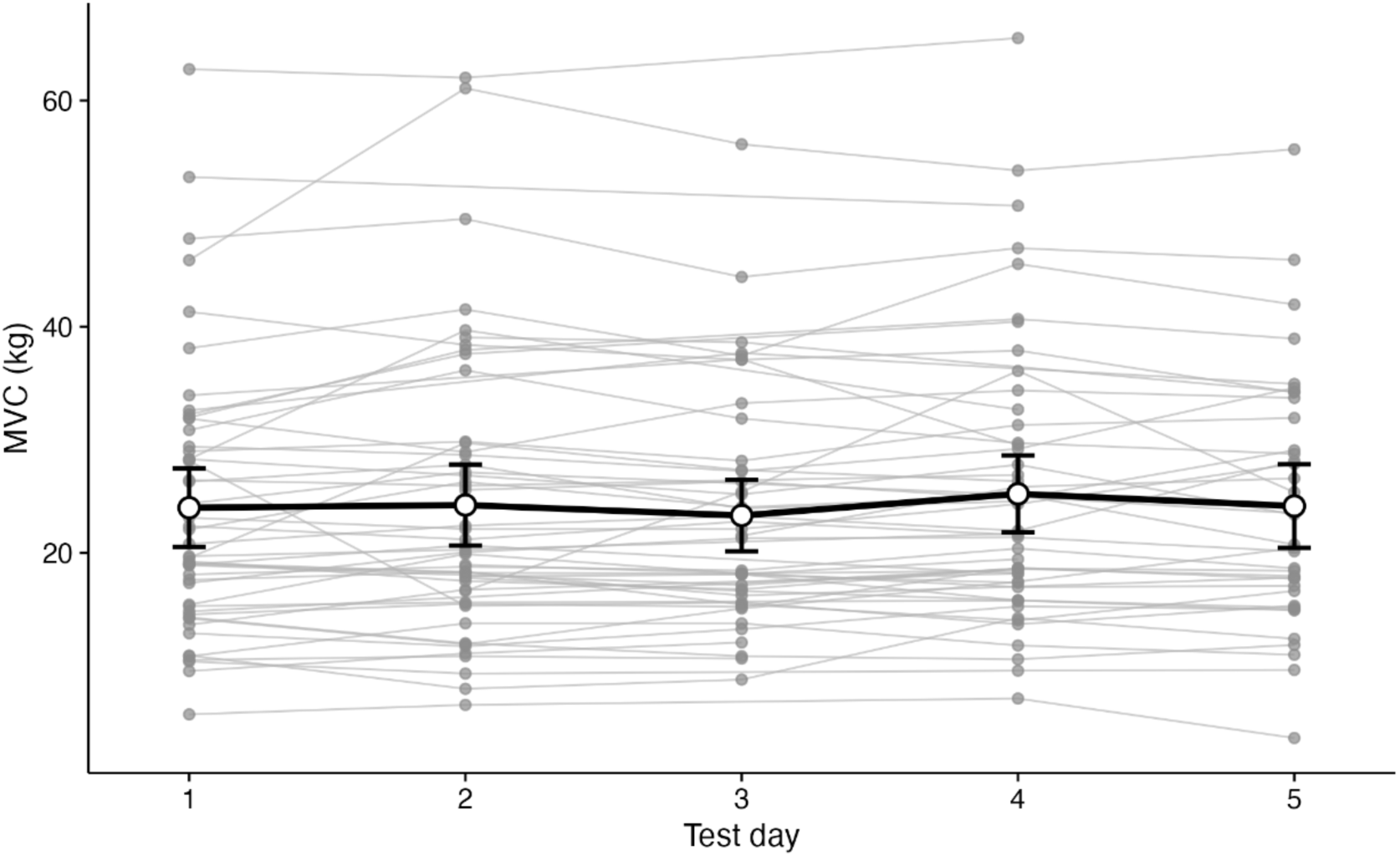
MVC across repeated laboratory test days. MVC force (N) was measured across repeated sessions conducted on separate laboratory test days within one week. Thin grey lines represent individual participant trajectories, whereas the black line and error bars indicate the group mean +/- SEM.

### Supplementary questionnaire phase comparisons

To assess the stability of questionnaire measures across testing sessions, late-follicular and mid-luteal scores were compared within the strict phase-verified sample. PHQ-9 scores did not differ between phases, with means of 8.63 (SD = 4.32) in the late-follicular phase and 8.72 (SD = 4.17) in the mid-luteal phase (*t*(42) = -0.17, FDR-adjusted *p* = 1.000, 95% CI [-1.188, 1.002]). GAD-7 scores were 6.98 (SD = 3.99) in the late-follicular phase and 5.81 (SD = 3.94) in the mid-luteal phase (*t*(42) = 2.18, FDR-adjusted *p* = .117, 95% CI [0.088, 2.237]). AMI scores were 25.12 (SD = 7.78) in the late-follicular phase and 26.65 (SD = 7.90) in the mid-luteal phase (*t*(42) = - 1.94, FDR-adjusted *p* = .117, 95% CI [-3.129, 0.059]). CIRENS scores were similar across phases at the group level, with means of -0.24 (SD = 1.56) and -0.24 (SD = 1.51), respectively (*t*(41) = 0.00, FDR-adjusted *p* = 1.000, 95% CI [-0.344, 0.344]).

### Supplementary decision-making descriptives, parameter recovery and additional moderator results

Descriptive performance metrics for the decision-making task are shown in Supplementary Fig. S2. Hard-choice proportions were similar across phases, with means of 53.72% (SD = 22.54%) in the late-follicular phase and 51.50% (SD = 19.59%) in the mid-luteal phase (*t*(43) = 1.28, *p* = .206, 95% CI [-1.27, 5.71]) (Fig. S2A). Task success was likewise similar across phases, with means of 89.46% (SD = 10.39%) and 88.79% (SD = 9.23%), respectively (*t*(43) = 0.38, *p* = .707, 95% CI [- 2.91, 4.26]) (Fig. S2B). Choice response times were also comparable across phases, with means of 2.89 s (SD = 0.86) and 3.05 s (SD = 1.48), respectively (*t*(43) = - 0.74, *p* = .466, 95% CI [-0.59, 0.28]). Catch-trial accuracy was high in both phases (late-follicular: 96.59%, SD = 12.75%; mid-luteal: 95.45%, SD = 14.54%).

Parameter recovery supported the adequacy of the computational model, with recovery correlations of *r* = .860 for effort sensitivity and *r* = .903 for reward sensitivity, and RMSE values of 2.542 and 3.405, respectively (Fig. S2C-D).

Other than mean-arousal and mean-valence effects reported in the main text, the remaining DM moderator models did not yield reliable phase-dependent effects. For effort sensitivity, neither DRSP nor HRV showed a significant interaction with phase, and the same was true for arousal variability and valence variability (all *p*s >= .118). For reward sensitivity, all tested phase interactions were non-significant, including DRSP, baseline HRV, active MVC HRV, HRV reactivity, mean arousal, arousal variability, mean valence, and valence variability (all *p*s >= .170). Thus, the robust DM moderator signal was specific to mean arousal and mean valence in relation to effort sensitivity, rather than reward sensitivity.

**Figure S2.**
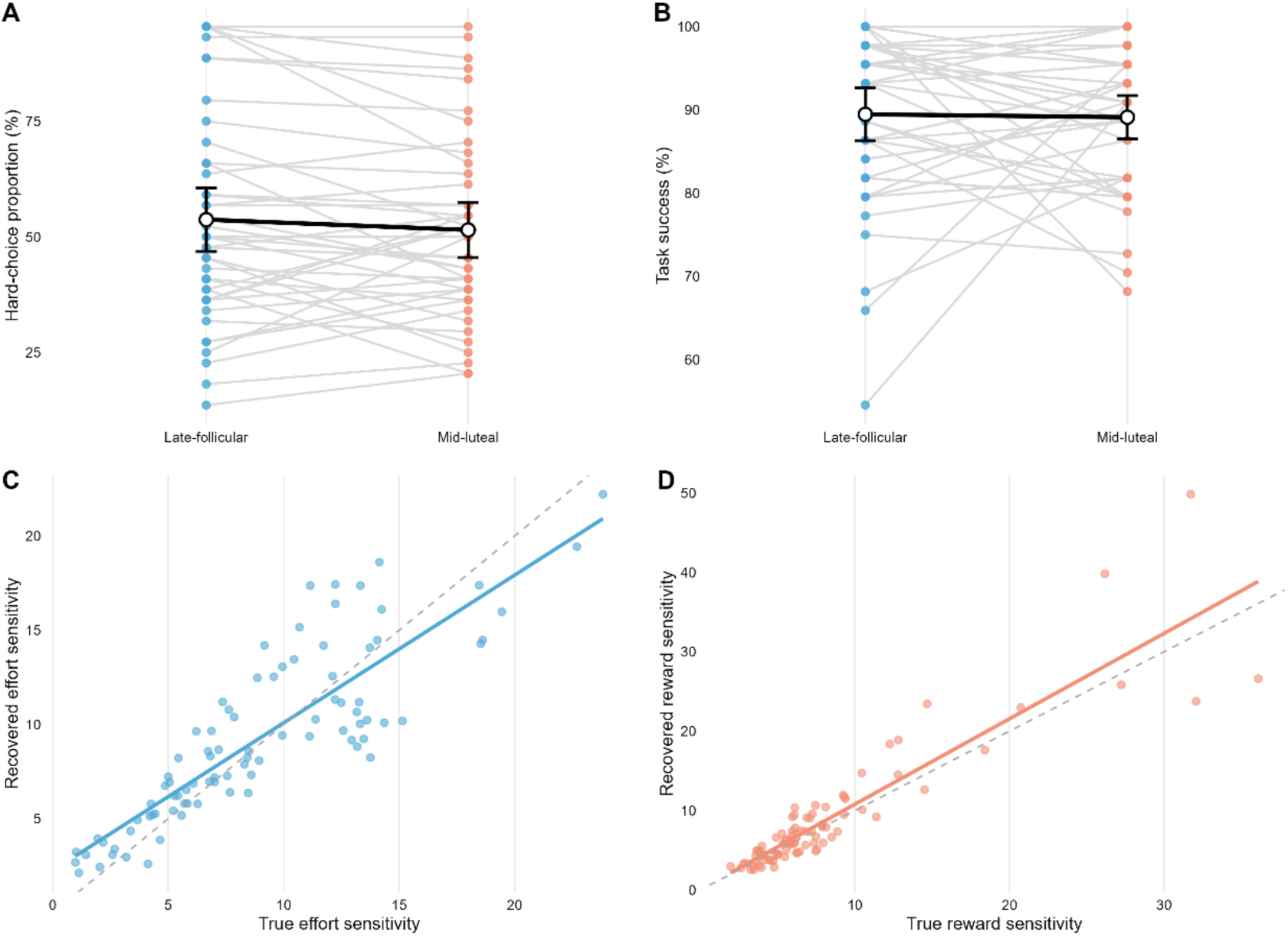
Supplementary decision-making descriptives and parameter recovery. (A) Hard-choice proportion (%) by phase. (B) Task success (%) by phase. In both panels, points show individual participants, connecting lines indicate within-participant pairing across phases, and black summary markers indicate phase means and 95% confidence intervals. (C) Parameter recovery for effort sensitivity. (D) Parameter recovery for reward sensitivity. Each point represents one simulated dataset, and the dashed diagonal indicates perfect recovery. Recovery correlations were *r* = .860 for effort sensitivity and *r* = .903 for reward sensitivity, and RMSE values of 2.542 and 3.405, respectively.

### Additional effort-perception moderator results

Arousal variability was not significantly associated with effort differentiation (beta = - 0.040, SE = 0.022, *t* = -1.79, *df* = 1148.87, *p* = .073, 95% CI [-0.083, 0.004]), and did not interact with phase (beta = -0.021, SE = 0.029, *t* = -0.72, *df* = 1148.87, *p* = .474, 95% CI [-0.078, 0.036]). Mean valence was likewise not associated with effort differentiation (beta = -0.007, SE = 0.018, *t* = -0.38, *df* = 1166.12, *p* = .706, 95% CI [-0.042, 0.028]), and did not interact with phase (beta = 0.035, SE = 0.028, *t* = 1.23, *df* = 1166.12, *p* = .217, 95% CI [-0.020, 0.090]). Thus, within the EMA analyses, the robust phase-dependent EP signal was specific to valence variability rather than to mean valence or arousal variability.

### Exploratory AMI analyses

In the EP task, higher AMI was associated with reduced effort differentiation (Fig. S3A). In the standardized model, the force x AMI interaction was significant (beta = - 0.039, SE = 0.013, *t* = -2.95, *df* = 1249.57, *p* = .003, 95% CI [-0.065, -0.013]), whereas the AMI main effect was not (beta = 0.048, SE = 0.029, *t* = 1.62, *df* = 196.05, *p* = .108, 95% CI [-0.010, 0.106]). In the DM task, higher AMI was associated with greater effort sensitivity (beta = 0.274, SE = 0.067, *t* = 4.06, *df* = 62.09, *p* < .001, 95% CI [0.139, 0.408]), but not with reward sensitivity (beta = - 0.150, SE = 0.098, *t* = -1.53, *df* = 84.22, *p* = .131, 95% CI [-0.345, 0.046]) (Fig. S3B).

**Figure S3.**
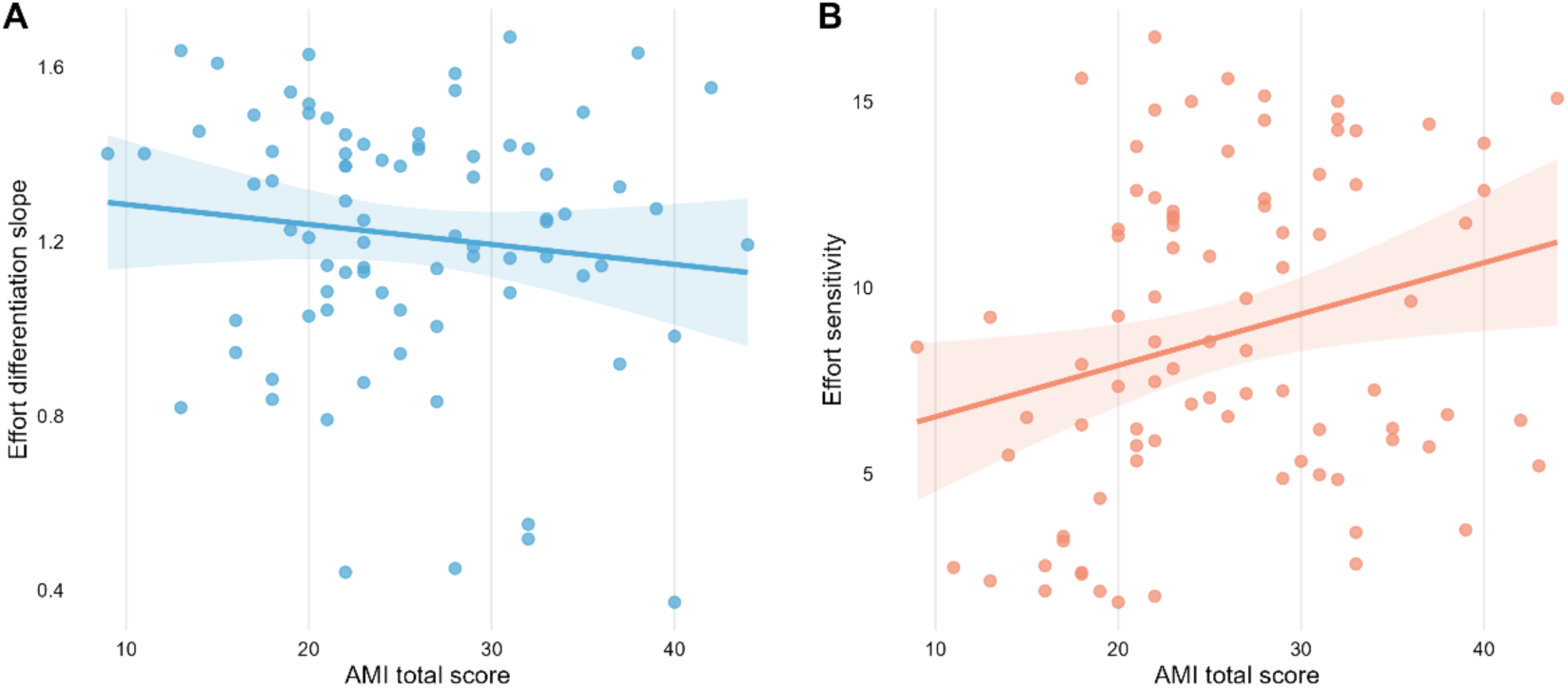
Exploratory associations between AMI and task-derived effort indices. (A) Association between AMI total score and effort differentiation slope in the effort-perception task. (B) Association between AMI total score and effort sensitivity in the decision-making task. In both panels, points represent one task session, lines show fitted linear trends, and shaded bands indicate 95% confidence intervals.

